# Development of RNA-based assay for rapid detection of SARS-CoV-2 in clinical samples

**DOI:** 10.1101/2020.06.30.172833

**Authors:** Vinod Kumar, Suman Mishra, Rajni Sharma, Jyotsana Agarwal, Ujjala Ghoshal, Tripti Khanna, Lokendra K. Sharma, Santosh Kumar Verma, Prabhakar Mishra, Swasti Tiwari

## Abstract

The ongoing spread of pandemic coronavirus disease (COVID-19) is caused by Severe Acute Respiratory Syndrome coronavirus 2 (SARS-CoV-2). In the lack of specific drugs or vaccines for SARS-CoV-2, demands rapid diagnosis and management are crucial for controlling the outbreak in the community. Here we report the development of the first rapid-colorimetric assay capable of detecting SARS-CoV-2 in the human nasopharyngeal RNA sample in less than 30 minutes. We utilized a nanomaterial-based optical sensing platform to detect RNA-dependent RNA polymerase (*RdRp*) gene of SARS-CoV-2, where the formation of oligo probe-target hybrid led to salt-induced aggregation and changes in gold-colloid color from pink to blue in visible range. Accordingly, we found a change in colloid color from pink to blue in assay containing nasopharyngeal RNA sample from the subject with clinically diagnosed COVID-19. The colloid retained pink color when the test includes samples from COVID-19 negative subjects or human papillomavirus (HPV) infected women. The results were validated using nasopharangeal RNA samples from suspected COVID-19 subjects (n=136). Using RT-PCR as gold standard, the assay was found to have 85.29% sensitivity and 94.12% specificity. The optimized method has detection limit as little as 0.5 ng of SARS-CoV-2 RNA. Overall, the developed assay rapidly detects SARS-CoV-2 RNA in clinical samples in a cost-effective manner and would be useful in pandemic management by facilitating mass screening.

## Introduction

Coronavirus disease is rapidly spreading across the world and raising severe global health concerns. In December 2019, China reported the first disease case in its Hubei Province. Based on the phylogenetic analysis, the identified novel coronavirus is named as Severe Acute Respiratory Syndrome Coronavirus 2 (SARS-CoV-2), and the disease spread by SARS-CoV-2 is known as “COVID-19”, declared as a pandemic by World Health Organization (WHO)(https://www.who.int/emergencies/diseases/novel-coronavirus-2019?gclid=EAIaIQobChMI-u-Pz9vW6QIVUyUrCh0kjAlOEAAYASAAEgJ0pvD_BwERemuzzi). Despite global massive efforts to control the outbreak of COVID-19, this pandemic is still on the rise. To date, lack of approved medicine or vaccine impede escalated the management of the COVID-19 epidemic. In the absence of an effective treatment strategy, developing affordable screening for rapid diagnosis is critically required in the management of COVID-19 (Udugama et al. 2020). According to the WHO, the immediate priority is the development of point-of-care tests for the detection of SARS-CoV-2 at an early stage with improved sensitivity(Report of the WHO-China Joint Mission on Coronavirus Disease 2019 (COVID-19); WHO 2017).

Currently, the COVID-19 diagnostic test falls into two categories: antibody and nucleic acid-based detection systems. The Developed immunoassay is rapid but inefficient for the detection of the pathogen at an early stage of infection.(Udugama et al. 2020) Besides, WHO does not currently recommend the use of antigen-based rapid diagnostic tests for patient care, but encourages the related research to improve their performance and potential diagnostic utility (Division of Viral Diseases. CDC 2019-Novel Coronavirus (2019-nCoV) Real-Time RT-PCR Diagnostic Panel; Division of Viral Diseases 2020).

Among the nucleic acid-based detection systems, the WHO considered a Real-time polymerase chain reaction (RT-PCR) based method as a gold standard for COVID-19 testing (Division of Viral Diseases. CDC 2019-Novel Coronavirus (2019-nCoV) Real-Time RT-PCR Diagnostic Panel; Division of Viral Diseases 2020). The sample-to-result time of the quantitative RT-PCR (qRT-PCR) was initially >4 hrs; however, constant efforts are underway to improve the turn-around time through automation. Besides, time-consuming process PCR based tests are expensive; they require sophisticated instruments and expertise (Udugama et al. 2020). Thus, other technologies, such as reverse transcription- loop-mediated isothermal amplification (RT-LAMP) (Kashir and Yaqinuddin 2020) and Genome-editing (Broughton et al. 2020), are being explored. These are promising technologies; however, the expected turn-around time would still be around 1 hour and may not be economical for mass screening, especially for the developing countries.

Nanomaterials based sensing platforms hold promise to develop rapid disease diagnostics. However, their limited use in the clinical setting is due to the need for sophisticated equipment (Chen et al. 2020; Qiu et al. 2020). A recent attempt to use antisense-nucleotide capped gold nanoparticles for N-gene based, COVID-19 detection could be a game changer (Moitra et al. 2020). Whereas, WHO suggested that N-gene has a relatively weak analytical capability, compared to the *RdRp* gene, to detect COVID-19 infection (Corman et al. 2020). Moreover, the authors have demonstrated COVID-19 detection in a cellular-model system, and application in human samples has yet to be shown.

In this study, we report for the first time gold nanoparticles (AuNPs) based rapid colorimetric assay for visual eye detection of COVID-19 RNA in human samples designed to target *RdRp* specific gene target in very cost-effective manner with wide application of mass screening in field. We utilized the surface plasmon resonance property of AuNPs to detect unamplified COVID-19 RNA in human samples. The ability of AuNPs to preferentially adsorb ssRNA/ssDNA over dsDNA/dsRNA is the crucial concept of this assay. The single and double-stranded oligonucleotides have different electrostatic properties, which provide stability (via retaining natural color) and aggregation (which causes color change) of AuNPs in solution, respectively (Jain et al. 2006).

## Materials and Methods

### Chemicals

Citrate buffer stabilized AuNPs (10 nm diameter) was purchased from Alfa Aesar (Thermo Fisher Scientific India Private Limited). Phosphate Buffer Saline (PBS pH 7.0) and NaCl were procured from Sigma Aldrich.

### Clinical samples

Nasopharyngeal RNA sample from subjects suspected have COVID-19 infectioon (n=136) were used. The COVID-19 testing of these subjects were done in the Indian Council of Medical Research (ICMR, India) approved diagnostic laboratory using Taqman-based RT-PCR kit (Labgun, lab, Genomics.co. Ltd, Republic of Korea). The study protocol to use clinical samples was approved by Institutional Human Ethics Committee SGPGIMS, Lucknow (Ref N. PGI/BE/327/2020). The cervical DNA samples from human papilloma virus (HPV) infected woman (n=2) was used as non-specific target control. The clinical Samples (cervical smear) for the HPV DNA test were processed using HPV Test Hybrid Capture® 2 protocol (QIAGEN). Samples with relative light units (RLU) /Cutoff Value ratios > 1.0 were considered as HPV positive and < 1.0 were considered as HPV negative. Only HPV positive DNA samples were used during validation experiments.

### In Vitro transcription (IVT) of SARS-CoV-2 RNA

The RNA (5ng) from confirmed COVID-19 positive human sample were first reverse transcribed with modified oligo dT primer having a T7 promoter sequence at its 5’ end, and the resulting single stranded cDNA was further *In-vitro* transcribed (IVT) using HiScribe T7 Quick High Yield RNA Synthesis Kit according to the manufacturer’s protocol (New England BioLabs Inc, NEB #E2050). The amplified RNA was than purified using Monarch RNA Cleanup Kit (New England BioLabs Inc.) and quantified by RNA HS reagent using Qubit system (Thermo Fisher Scientific).

### Colorimetric assay for the detection of SARS-CoV-2 RNA

For colorimetric assay, reaction was set up in 10 μL of reaction volume in sterile PCR tubes, containing **(a)** hybridization buffer containing 80mM NaCl), **(b)** 0.5 μM oligo probe for *RdRp* genes, and **(c)** target RNA from positive patients or IVT RNA. The *RdRp* oligo probe (5’-GTGATATGGTCATGTGTGGCGG-3’) was used to specifically detect the presence of SARS-CoV-2 RNA in the assay. In parallel, to measure the specificity of the assay, input genomes from different source such nasopharyngeal RNA from COVID negative subjects and HPV DNA from cervical cancer positive samples were used as negative controls. Similarly, non-template control (NTC) was also included to measure the background reactivity. In addition, to confirm the working principle of the assay, a different RNA template (isolated from pancreas) and pancreas specific *REG-3* (Regenerating islet-derived protein 3) gene oligo probe (5’-GTGCCTATGGCTCCTATTGCT-3’) were used separately.

The final reaction mixture with above combinations was then denatured at 95°C for 30 seconds, annealed at 60°C for 60 seconds and then cooled to room temperature for 10 minutes. Subsequently, 10 nM colloidal AuNPs (~10 nm) were added to the assay mixture and allowed to develop color for 1-2 minute.

### Spectral studies and measurement of sensitivity

Absorption spectrum of the assay mixture was recorded in the range of 300 -700 nm. The peak shift from 520 nm (known as red-shift) and peak broadening after 520 nm were measured as a characteristic feature of the salt induced aggregation. Using various combinations of positive and negative controls (as discussed in the previous section) the specificity of reaction and aggregation were compared. Assay sensitivity was determined by serially diluting the input SARS-CoV-2 RNA from both the IVT synthesized and, synthetic SARS-CoV-2 control (nCov19 control kit by Applied Biosystems) ranging from 5-0.1ng and 1-0.1ng concentrations respectively.

### Statistical Analysis

Diagnostic accuracy of the new colour test was calculated by assuming RT-PCR as gold standard method. Sensitivity (True Positive rate), Specificity (True Negative rate) and overall accuracy (True positive and true negative rate) were calculated with 95% confidence interval. Likelihood ratio positive (sensitivity / false positive rate), Likelihood ratio negative (false negative rate / specificity), Positive predictive value (True positive value / Total positive results predicted by colour test), Negative predictive value (True negative value / Total negativeresults predicted by colour test) were also calculated. Measured accuracy was considered statistically significant (p<0.05) when 50% did not falling within the confidence limit for values given in % whereas LR are considered significant when 1 was falling within confidence limit. Statistical analyses were performed using software MedCalc for Windows (MedCalc Software, Ostend, Belgium).

## Result and discussion

In this study, we report for the first time the development of a rapid and affordable RNA-based assay for the visual detection of the SARS-CoV-2 genome in human samples. Using RT-PCR as gold standard, the developed assay was found to have a sensitivity of 85.29% and sepcififcity of 94.12%. For the assay, we use surface plasmon resonance property of gold nanoparticles/colloids (AuNP) and targeted the *RdRp* specific gene sequence of SARS-CoV-2. *RdRp* is essential for viral replication and has higher analytical power than E (envelope protein) and N (nucleocapsid protein) genes of SARS-CoV-2 (Corman et al. 2020). Our current established assay, salt-induced aggregation and color change of the gold colloids occurs after *RdRp* oligo probe hybridizes with its specific target RNA of SARS-CoV-2. With the current escalated demand of cost effective, easy and sensitive diagnostic for COVID-19, the test was developed using commercially available nCoV19 synthetic DNA and validated further using clinical samples from COVID-19 subjects (as confirmed using Taqman based RT-PCR method, Table S1). In our study, we demonstrated a visual change in gold colloid color from pink to blue when RNA samples from subjects with clinically diagnosed COVID-19 infection hybridize with *RdRp* oligo probe. Simultaneously, the color remained pink in SARS-CoV-2 negative samples due to the absence of hybridization.

In the study, out of 136 samples, 50% samples (n=68) were true positive (COVID-19 Positive) and 50% samples (n=68) true negative (Free from COVID-19 disease) confirmed by gold standard diagnostic test RT-PCR. These true positive and negative samples were again tested by New Colour test to assess the diagnostic accuracy of our new test **(Table 1).** Result showed that sensitivity (true positive rate) of the new test was 85.29% (58/68) and specificity (true negative rate) of 94.12% (64/68) with statistically significant (each p value<0.05). Overall diagnostic accuracy of the new colour test was 89.71% (122/136) with statistically significant (p<0.05). Likelihood ratio positive indicated that new colour test has 14.50 times more chances to detect true positive rate (w.r.t. false positive rate) and Likelihood ratio negative of 0.16 i.enew colour test has 6.25 times more chances to detect true negative rate (w.r.t. false negative rate).(each p<0.05). Positive predicted value (PPV) of 93.55% showed that probability to the correct prediction of true positive results from its own positive predicted results would be 93.55% and Negative predictive value (NPV) of 86.49% showed that probability to the correct prediction of true negative results from its own negative predicted results would be 86.49% (each p<0.05) **(Table 2).**

**Table 1:** Comparison of New Colour test by comparison to RT-PCR. (N=136)

| Table 1: Comparison of New Colour test by comparison to RT-PCR. (N=136) |  |  |  |  |
| --- | --- | --- | --- | --- |
|  |  | **RT-PCR**<br>**(Gold Standard Method)** |  | Total |
|  |  | Positive | Negative |  |
| **New Colour**<br>**Test** | Positive | 58 | 4 | **62** |
|  | Negative | 10 | 64 | **74** |
|  | **Total** | **68** | **68** | **136** |

**Table 2:** Diagnostic Accuracy of New Colour test by comparison to RT-PCR. (N=136)

|  |  |  |  |  |
| --- | --- | --- | --- | --- |
| Measurements | Value | 95% CI |  | P Value |
|  |  | Lower | Upper |  |
| Sensitivity<br>(True positive rate) | 85.29% | 74.61% | 92.72% | <0.05 |
| Specificity<br>(True negative rate) | 94.12% | 85.62% | 98.37% | <0.05 |
| Likelihood Ratio Positive<br>(LR+) | 14.50 | 5.58 | 37.71 | <0.05 |
| Likelihood Ratio Negative<br>(LR-) | 0.16 | 0.09 | 0.28 | <0.05 |
| Positive Predictive Value<br>(PPV) | 93.55% | 84.79% | 97.42% | <0.05 |
| Negative Predictive Value<br>(NPV) | 86.49% | 78.26% | 91.92% | <0.05 |
| Overall Accuracy | 89.71% | 83.33% | 94.26% | <0.05 |
| Above estimation are based on equal number of disease- and disease-free cases in study samples detected by Gold standard method. |  |  |  |  |

Figure 1 illustrates the sequential schematics of process of the developed test. The color of the gold colloid solution is dependent on the aggregation property of AuNPs in suspension (Hunter 1991; Lazarides and Schatz 2000). The aggregation-induced color change can be visually monitored (by the naked eye) and quantitated through absorption spectroscopy. Generally, in aqueous solution, gold colloids remain stabilized by the coating of negatively charged citrate ions (Grabar et al. 1995) and have visible appearance of pink color (Figure 1). In solution, individual particle exhibits a surface plasmon resonance peak (λmax) at 520 nm (Figure 2a, Green curve). The oligonucleotide probe preferentially adsorbs on AuNPs and provides additional stability due to the addition of negative charges (Li and Rothberg 2004). The same phenomenon was observed in our experiments when *RdRp* oligo probe adsorbed and protected the salt-induced aggregation of colloid in the absence of target RNA (NTC) (Figure 2a, Grey curve). Except for the reduced intensity of the absorption spectrum, which was due to dilution of the colloid (Figure 2a, Grey curve), NTC and AuNP assays showed no change? A pure colloid is pink in color (Inset of Figure 2a left vial), which turns into light pink when it reacts with hybridization buffer containing oligo probe in equal volume (without target) (Inset of Figure 2a right vial). Unlike dsDNA, inherent structural flexibility of ssDNA/RNA to partially uncoil its bases, exposing them to AuNPs, generates the attractive electrostatic forces causing them to allow over colloids and giving protection against electrostatic interaction causing salt-induced aggregation (Li and Rothberg 2004).

**Figure 1.**
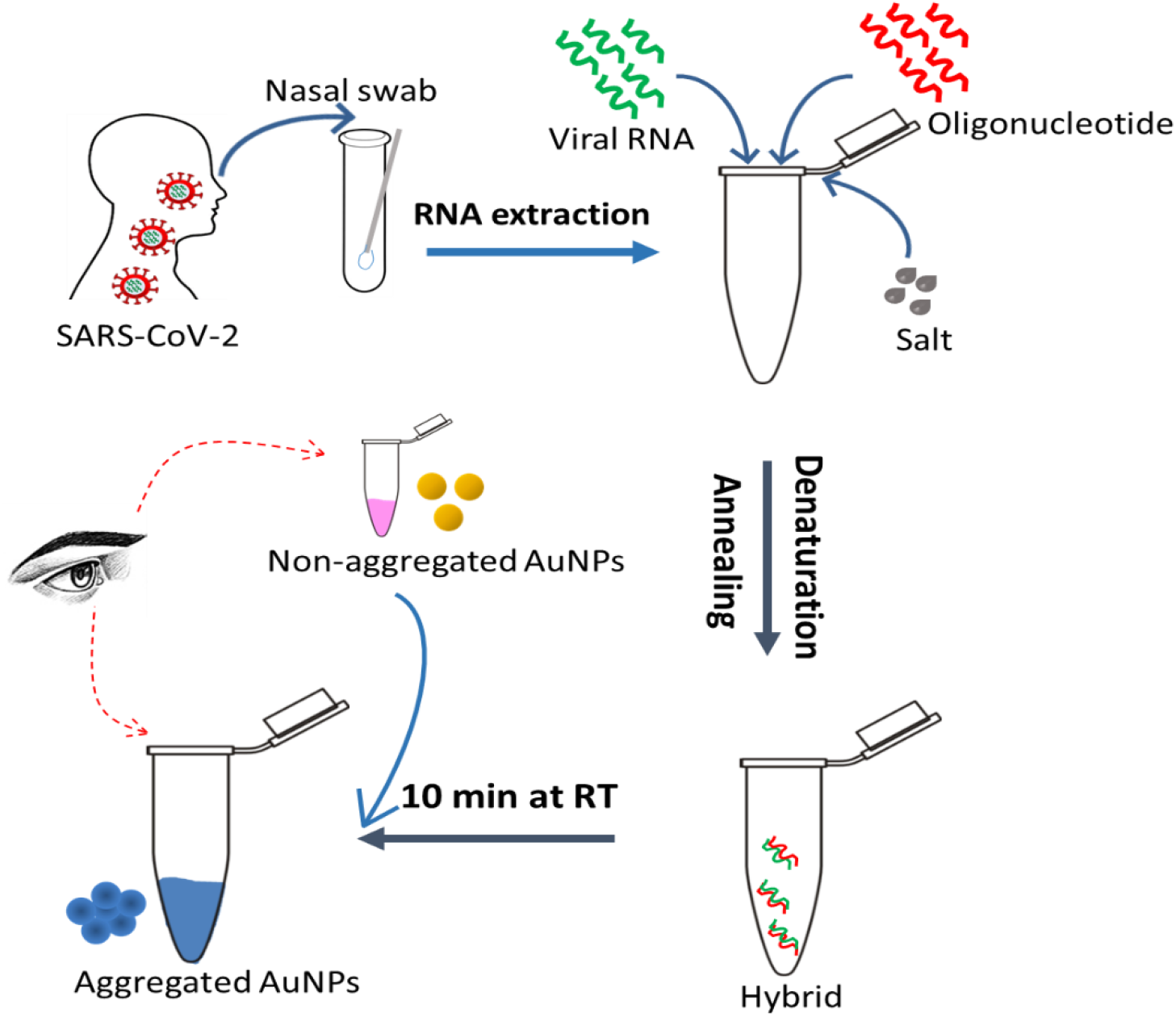
Schematic representation of the assay for the visual detection of SARS-CoV-2 RNA. Schematic illustrates the assay flow to detect *RdRp* (RNA dependent RNA polymerase) gene sequence of SARS-CoV-2 in the nasopharyngeal RNA sample from subject clinically diagnosed with nCOVID infection (**positive control**). Hybridization buffer with *RdRp* oligo probe (forward) was mixed with RNA sample. The reaction mixture was denatured at 95°C for 30 seconds, followed by annealing at 60° C for 60 seconds. After annealing, the tube was kept at room temperature for 10 minutes before colloidal AuNPs were added. A pure colloid is pink in color. It turns blue in the vial containing RNA sample from **positive control** due to salt-induced aggregation upon successful hybridization between oligo probe and target RNA. The color remained pink in the absence of target RNA, or the presence of a non-specific target.

**Figure 2.**
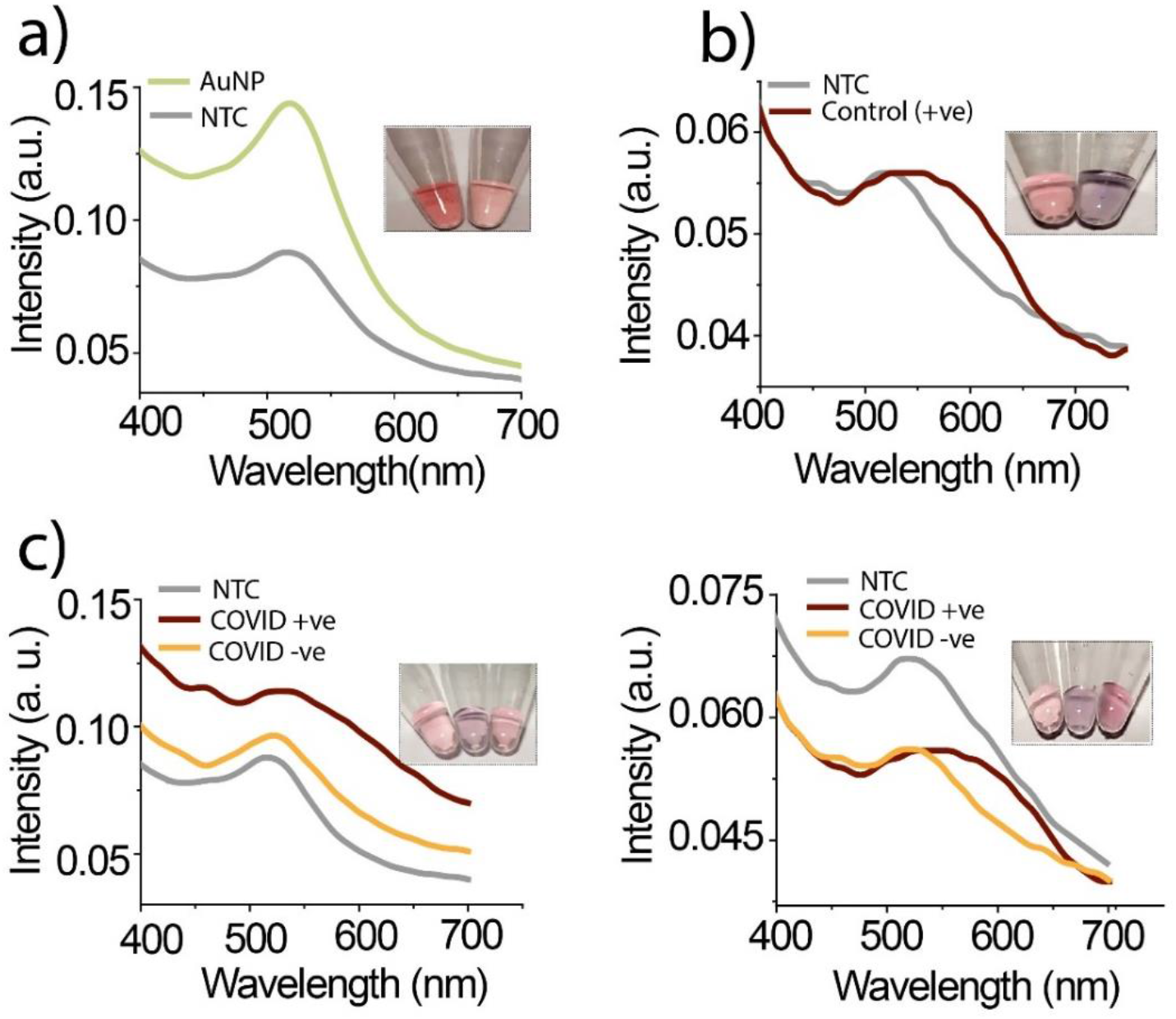
Colorimetric assay to detect SARS-CoV-2 RNA. **(a)** Comparative absorption spectra of unmodified AuNPs (Green curve) and oligo probe stabilized AuNPs i.e., NTC (Grey curve). Both spectra are exhibiting characteristics absorption peak at λmax 520 nm, however, the reduced peak intensity in NTC is due to dilution of colloid solution. In NTC, no red-shift in peak position, approve the stabilizing property of single-stranded oligo probe, Optical images of gold colloids (left vial) and NTC (right vial) are shown in the inset. **(b)** Comparative absorption spectra of NTC (Grey curve) and positive control i.e., nasopharyngeal RNA sample from subject clinically diagnosed with nCOVID infection (Brown curve). Broadening of the peak as well as red-shift in peak position confirm the salt-induced aggregation of AuNP due to successful hybrid formation in control. Optical images shown in inset demonstrate the evident change in the color of the solution from pink to blue in the control vial (right) while no change in color of NTC vial (left). **(c)** representative absorption spectra, and in the inset shows optical images comparing assay performed with NTC (Grey curves, left vial), RNA from clinically diagnosed nCOVID infected subjects (Brown curve, middle vial), and RNA from subjects without nCOVID infection (Yellow curve, right vial). Samples from a total of eighteen infected and eighteen uninfected individuals were analyzed (optical images were attached as supplimentry figure S2).

In this assay, SARS-CoV-2 RNA from human patients or IVT synthesized RNA was added into hybridization buffer (containing oligo probe), followed by denaturation and annealing at 95°C (30s) and 60°C (60s), respectively. After cooling at room temperature, the gold colloid was added into the above reaction mixture. The colloid color changes in visible range from pink to blue, indicate the formation of hybridized product (Figure 2b-c, and Figure 3). Broadening of the peak with red-shift (~30 nm) was observed in the spectrum of aggregated colloids than non-aggregated, confirms the success of developed assay for detection of an unamplified target with unmodified colloids in a quick and facile way. The principle of binding oligo probe to its specific target leading to change in color of the solution was independently verified using a different template RNA (isolated from pancreas tissue) and pancreas specific gene *REG3* oligo probe in a separate assay. This assay also resulted in a similar change in color and absorption spectra as optimized earlier for SARS-CoV-2 RNA and *RdRp* oligo probe. It established the working principle and specificity of the test (Figure S1).

**Figure 3.**
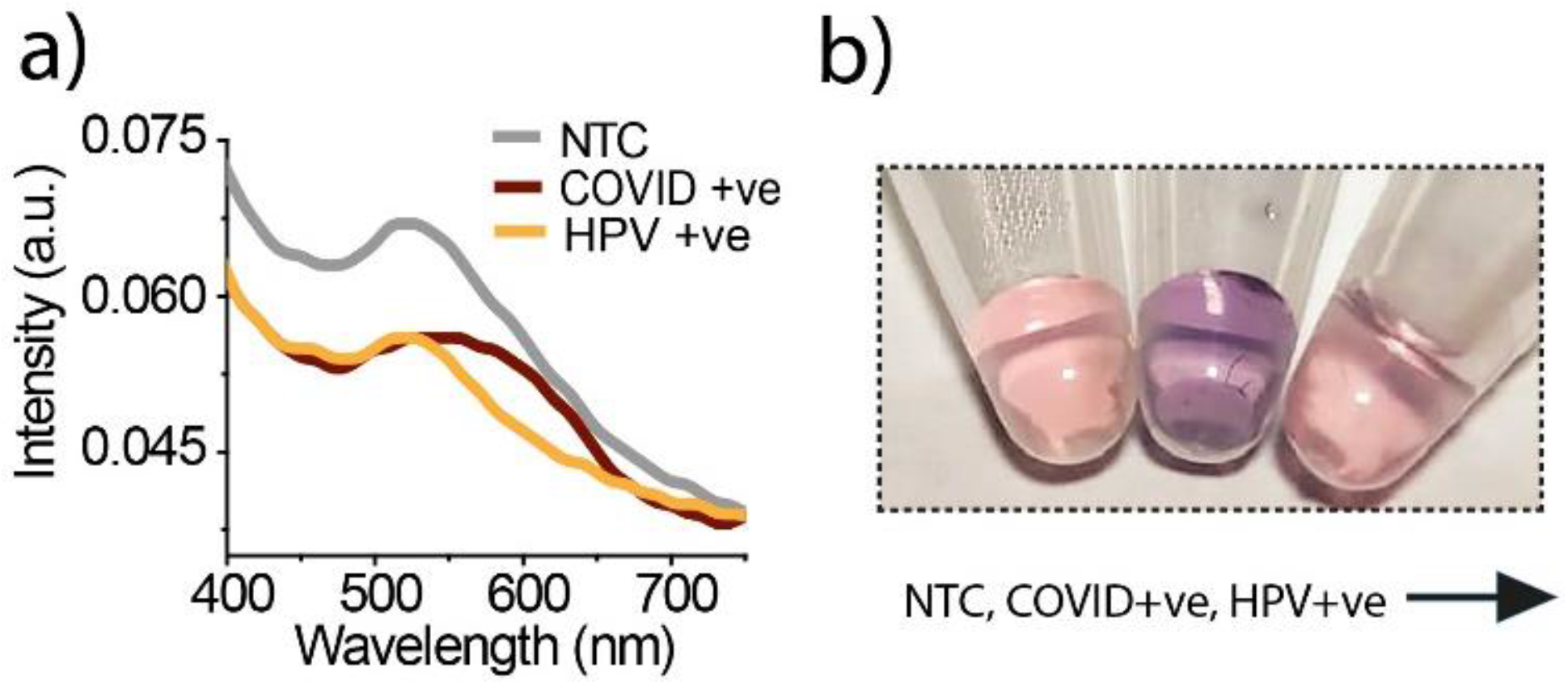
Specificity of developed assay to detect SARS-CoV-2 RNA. Nasopharyngeal-RNA from nCOVID infected subject (positive control), and a cervical DNA sample from Human papillomavirus (HPV, non-specific target control) infected women were tested. **(a)** Comparative absorption spectra for no target control (NTC, Grey curve), positive control (Brown curve), and non-specific target control (HPV, Yellow curve). In the positive control, the broadening of the peak and red-shift in peak position (Brown curve) confirmed the salt-induced aggregation upon successful hybrid formation. **(b)** Optical pictures demonstrate that the colloid color from pink to blue changes only in the vial with the positive control (middle vial), while the color remained pink in the vial with NTC (left vial) or HPV (right vial).

We determined the cross-reactivity using a cervical-DNA sample from women diagnosed with HPV infection (non-specific target control). No color change of gold colloids was observed with HPV DNA, indicating no hybridization, and specificity of the developed assay (Figure 3 b, right vial). Contrary to HPV DNA-negative control and NTC, a positive control sample shows development of blue color (Figure 3 b, middle vial). Absorption spectrum (of colloids) with HPV DNA-negative control exhibited characteristics similar to that of NTC, and no red-shift or peak broadening as found with positive samples (Figure 3a). Cross-reactivity of the developed assay with other respiratory viruses is warranted. However, we do not anticipate the same as the test utilizes the detection of *the RdRp* gene of the SARS-CoV-2 virus. The oligo probe sequence used in our assay is not complementary to any human mRNAs and other members of the SARS family, as verified by BLAST using the NCBI database.

IVT synthesized SARS-CoV-2 RNA was used to test the sensitivity of the developed assay. RNA ranging from 0.1 to 5ng resulted in a gradual change in colloid color from light pink to blue (Figure 4b). The color difference at 0.1 ng, compared to NTC, is barely visible with naked eyes (Figure 4 b extreme left vial). However, the absorption spectra of colloids show a clear red-shift with peak broadening up till 0.5 ng target RNA (Figure 4a).

**Figure 4.**
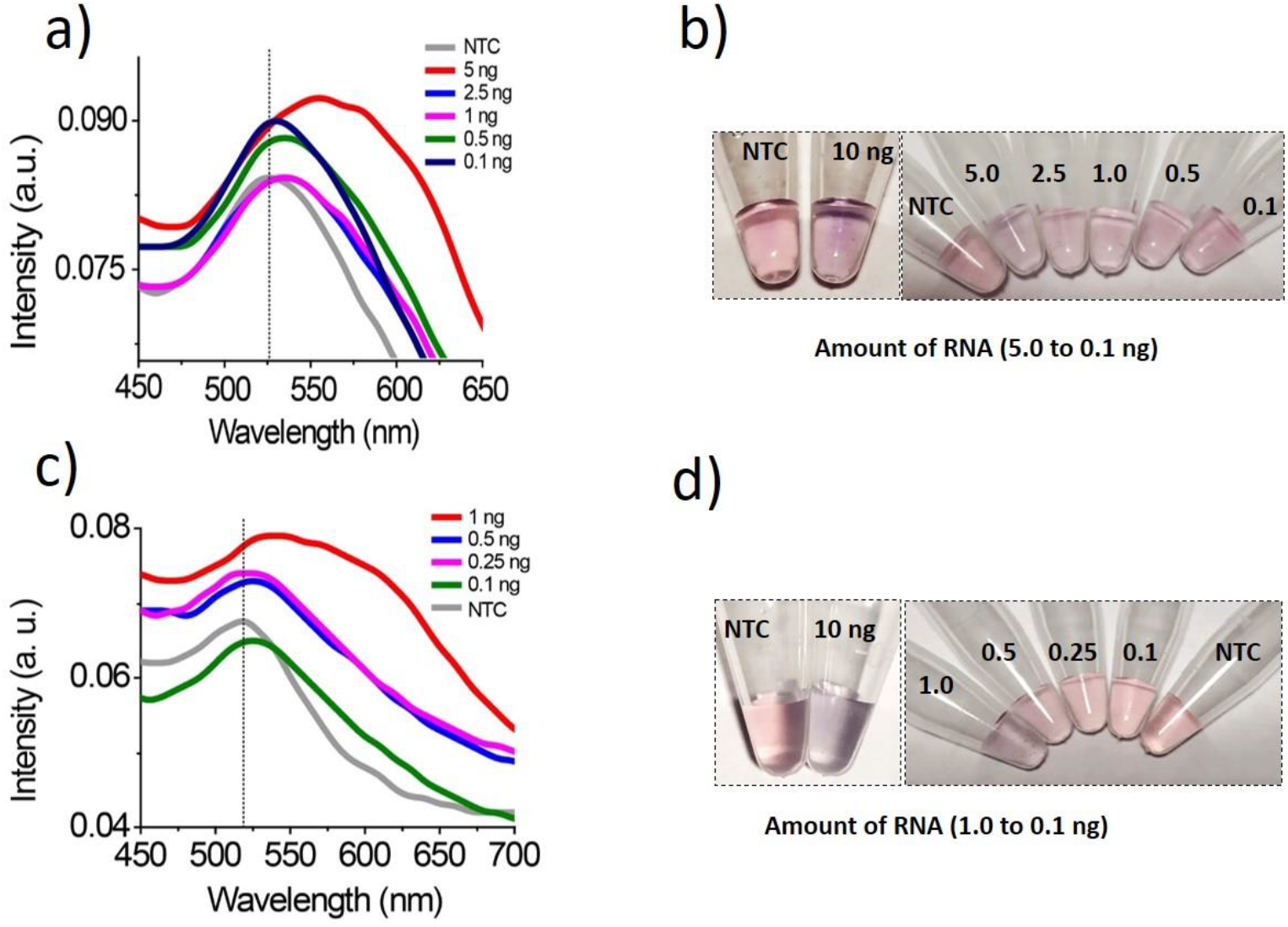
Sensitivity of developed assay to detect SARS-CoV-2 RNA. Assay sensitivity was determined using different concentrations (concentration are in ng) of IVT synthesized SARS-CoV-2 RNA **(a-b)**, or nCOVID synthetic DNA **(c-d)**. Absorption spectra **(a and c)** corresponding to aggregated colloids exhibit a clear red-shift in peak with broadening, indicating successful hybridization. Optical pictures **(b &d)** of the assay performed to demonstrate the color change, an extra pair of vials (on the left) to show color change with a higher amount (10ng) of target nucleic acid for the reference

We also determined the assay sensitivity using different dilutions (1-0.1ng) of PCR-amplified synthetic DNA (positive control, with nCoV19 control kit). Similar to IVT synthesized RNA, a decreasing amount of DNA (from 1 to 0.1 ng) show a gradual change in colloid color from blue to light pink (left to right, Fig. 4d). A clear red-shift with peak broadening reflects in the absorption spectra of colloids recorded with control DNA, compared to NTC (Fig. 4c). Accordingly, positive control showed a clear visual demarcation up to 0.5 ng amount, compared to NTC (Fig. 4d). A similar recent approach utilizes thiol capped gold nanoparticles to detect N-gene of the SARS-CoV-2 gene in a cellular system (Moitra et al. 2020). However, for the detection of SARS-CoV-2 RNA in human samples, the N-gene has reportedly inferior analytical power than the *RdRp* gene (Corman et al. 2020). Thus, it would be essential to know the assay's performance, developed by Moitra et al., with clinical samples.

## Conclusions

We have successfully developed an affordable gold nanoparticles-based colorimetric test for the rapid detection of SARS-CoV-2 RNA in humans. The assay can detect up to 0.5 ng of SARS-CoV-2 RNA. The turnaround time of our assay is less than 30 minutes. Moreover, the developed test will be helpful for mass screening, as it does not require sophisticated equipment. However, while analyzing the clinical samples we observed the assay works best with the freshly isolated RNA samples. In addition, pH and salt concentration in the elution buffer, used for RNA isolation, may affect the result and hence optimization may be needed when using a different kit for RNA isolation.

## Acknowledgement

The study was supported by intramural grants (A-24-PGI/IMP/81/2020) and overhead funds from the extramural grants to ST from DBT, ICMR and MHRD. The authors wish to thank the technical staff of the Department of Microbiology (SGPGIMS, Lucknow and RMLIMS, Lucknow) for RNA extraction and RT-PCR analysis for the clinical diagnosis of subjects.

## Contributions

ST, VK; conceived the idea; ST, VK, SM, RS; performed the experiments, JY, UG; provided clinical samples and the diagnosis; TK, LS, PM and SKV; provided critical comments on the manuscript draft; ST, VK and SM; drafted the manuscript.

## Supplementary Files

Supplementary information supporting the finding of this study is available in this article as supporting information.

## Supporting information

**Figure S1.**
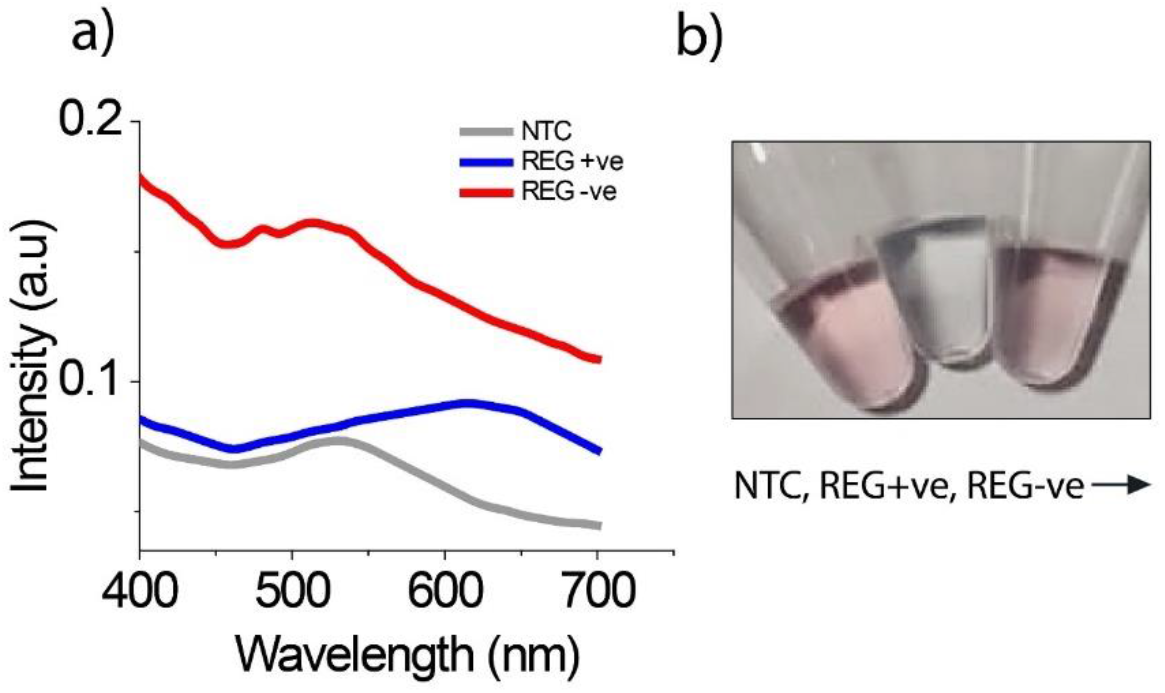
The assay was performed using an oligo probe specific for regenerating islet-derived (REG 3). The assay was containing RNA extracted from pancreatic tissue (positive control), placental RNA (negative control), or no RNA (NTC). **(a)** Comparative absorption spectra of NTC (Grey curve), positive control (Blue curve), and negative control (red curve). In the positive control, the broadening of the peak and red-shift in peak position (Blue curve) confirm the salt-induced aggregation upon successful hybrid formation. **(b)** Optical pictures demonstrate that the colloid color from pink to blue changes only in the vial with the positive control (middle vial), while the color remained pink in the vial with NTC (left vial) or negative control (non-specific target, right vial).

**Figure S2.**
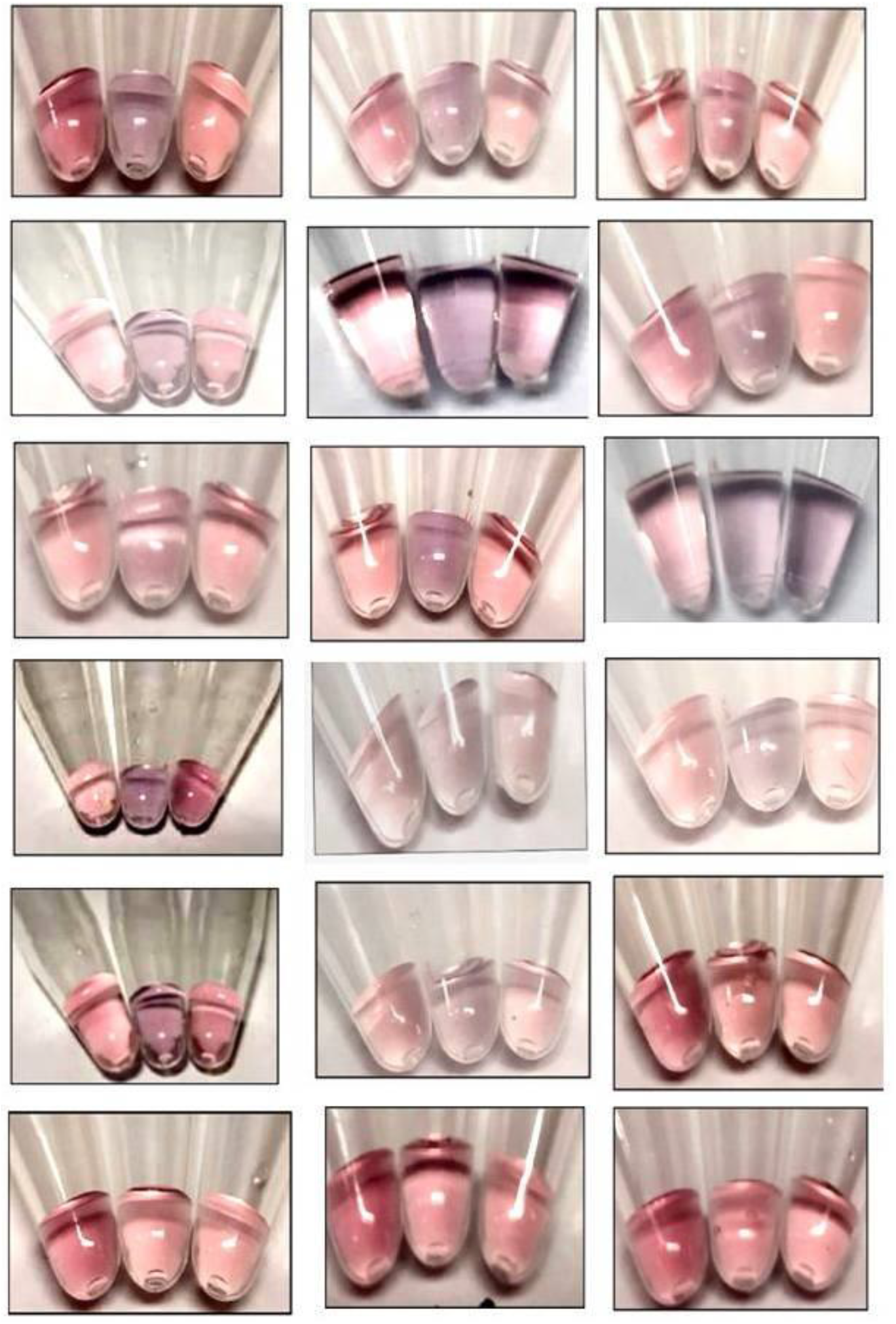
*The figure* shows optical images comparing assay performed with NTC (left vial), RNA from clinically diagnosed nCOVID infected subjects (middle vial), and RNA from subjects without nCOVID infection (right vial). *Clinical diagnosis of the subjects based on Taqman RT-PCR analysis is given in Table S1*

**Table S1.**
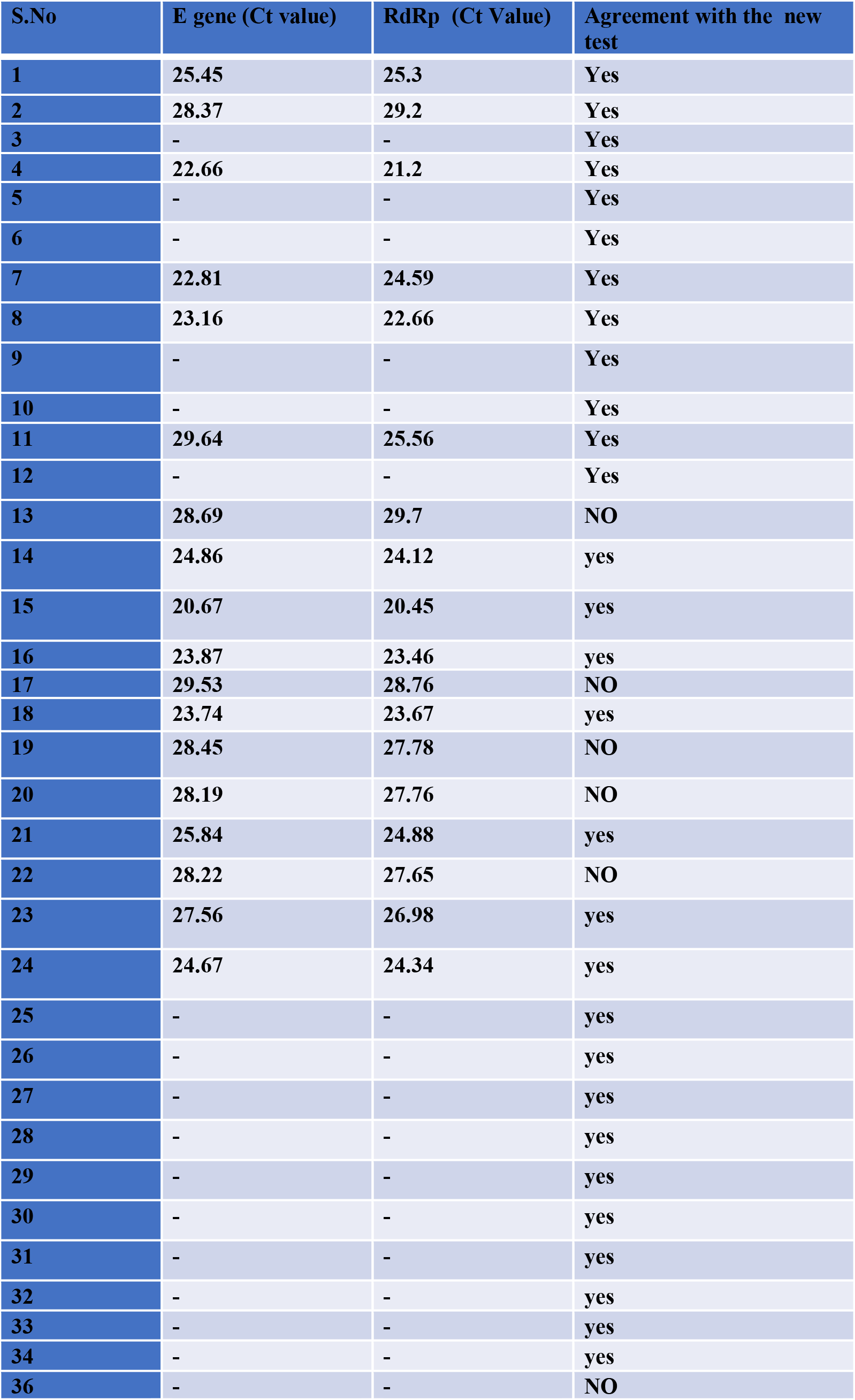
The table shows the clinical diagnosis of the subjects based on Taqman RT-PCR analysis. RNA from the nasopharyngeal samples were analyzed by the new test (optical images shown in Figure S2). RNA samples received from clinical laboratories were re-elluted in RNAase free water before testing with new method.

**Figure S3.**
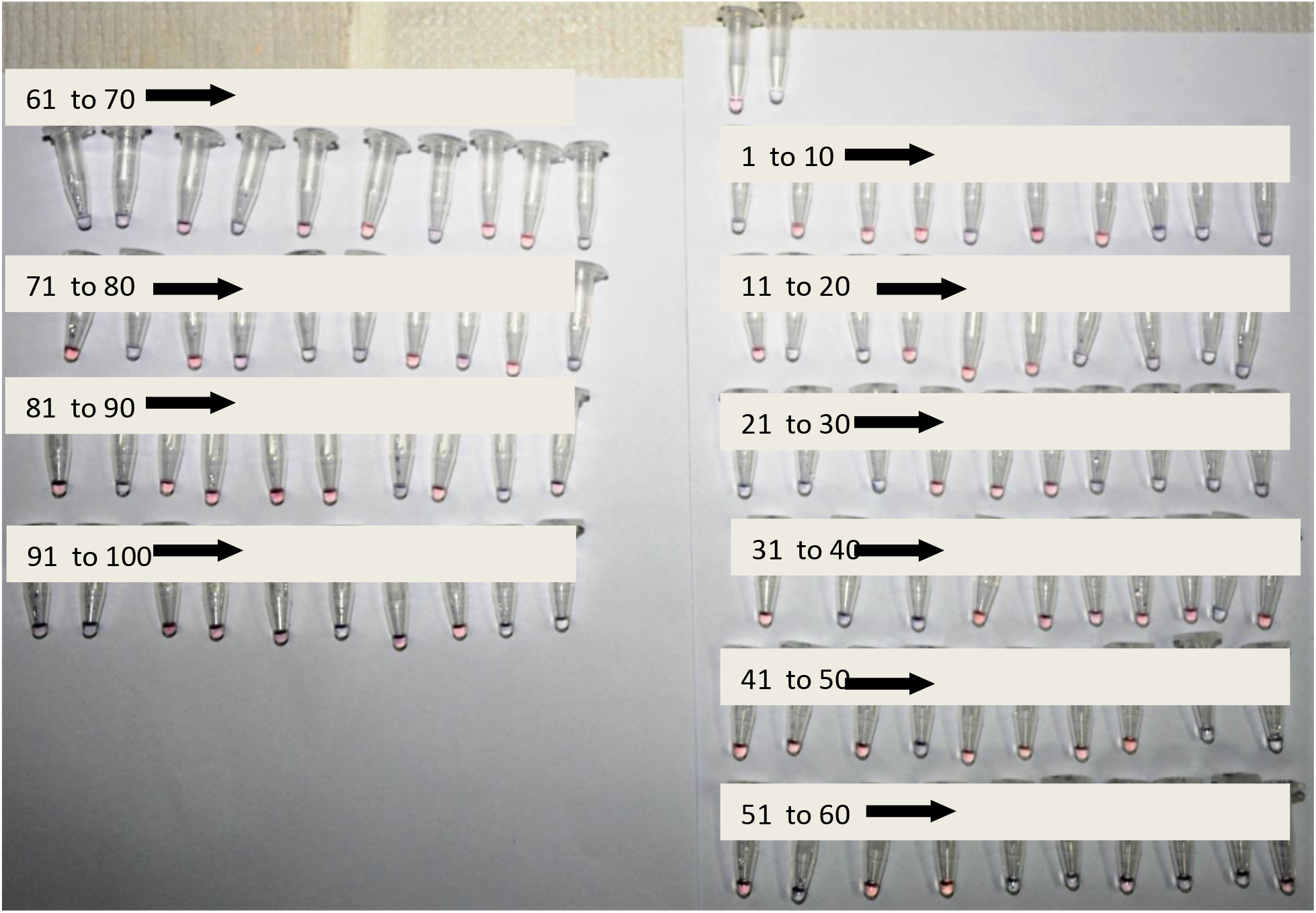
*The figure* shows optical images of the assay performed using 100 clinical samples from clinical laboratory at RMLS (Lucknow). *Clinical diagnosis of the subjects based on Taqman RT-PCR analysis is given in Table S2*

**Table S2.** The table shows the clinical diagnosis of the subjects based on Taqman RT-PCR analysis. RNA extracted from the nasopharyngeal samples were analyzed by the new test (optical images shown in Figure S3). Nasopharyngeal samples Samples were collected and RNA was isolated at RMLMS (Lucknow).

| Si No. | Code | Age | Gender | RT-PCR result (RMLMS) | E-Gene (CT value) | N- Gene (CT value) | RdRp (Ct value) | New test* |
| --- | --- | --- | --- | --- | --- | --- | --- | --- |
| 1. | 1 | 39 | M | POSITIVE | 23.14 | 24.37 | 23.64 | Yes |
| 2. | 2 | 47 | M | NEGATIVE | - | - |  | Yes |
| 3. | 3 | 57 | M | NEGATIVE | - | - |  | Yes |
| 4. | 4 | 53 | M | NEGATIVE | - | - |  | Yes |
| 5. | 5 | 48 | F | POSITIVE | 26.67 | 26.44 | 26.21 | Yes |
| 6. | 6 | 62 | F | NEGATIVE | - | - |  | Yes |
| 7. | 7 | 38 | F | NEGATIVE | - | - |  | Yes |
| 8. | 8 | 43 | M | NEGATIVE | - | - |  | **NO** |
| 9. | 9 | 62 | F | POSITIVE | 21.17 | 21.36 | 21.68 | Yes |
| 10. | 10 | 57 | M | POSITIVE | 18.93 | 19.43 | 19.13 | Yes |
| 11. | 11 | 36 | F | NEGATIVE | - | - |  | Yes |
| 12. | 12 | 48 | M | POSITIVE | 29.04 | 29.46 | 29.44 | Yes |
| 13. | 13 | 48 | F | POSITIVE | 18.15 | 19.34 | 19.65 | Yes |
| 14. | 14 | 60 | M | NEGATIVE | - | - |  | Yes |
| 15. | 15 | 50 | F | NEGATIVE | - | - |  | Yes |
| 16. | 16 | 45 | M | POSITIVE | 22.64 | 23.68 | 21.92 | **NO** |
| 17. | 17 | 47 | M | POSITIVE | 27.78 | 27.16 | 27.04 | Yes |
| 18. | 18 | 49 | F | POSITIVE | 28.79 | 28.41 | 27.86 | Yes |
| 19. | 19 | 39 | F | POSITIVE | 29.56 | 28.18 | 29.03 | Yes |
| 20. | 20 | 71 | M | POSITIVE | 27.56 | 27.86 | 27.13 | Yes |
| 21. | 21 | 72 | M | POSITIVE | 28.92 | 28.48 | 27.89 | Yes |
| 22. | 22 | 58 | F | POSITIVE | 30.04 | 29.67 | 29.16 | Yes |
| 23. | 23 | 61 | F | POSITIVE | 30.86 | 30.64 | 30.78 | Yes |
| 24. | 24 | 59 | F | NEGATIVE | - | - |  | Yes |
| 25. | 25 | 52 | M | NEGATIVE | - | - |  | Yes |
| 26. | 26 | 51 | M | NEGATIVE | - | - |  | Yes |
| 27. | 27 | 50 | M | POSITIVE | 29.06 | 28.06 | 29.65 | Yes |
| 28. | 28 | 40 | F | POSITIVE | 31.27 | 30.56 | 29.85 | Yes |
| 29. | 29 | 35 | M | POSITIVE | 28.06 | 28.67 | 28.96 | Yes |
| 30. | 30 | 28 | F | POSITIVE | 24.06 | 24.56 | 23.94 | Yes |
| 31. | 31 | 30 | M | NEGATIVE | - | - |  | Yes |
| 32. | 32 | 37 | F | POSITIVE | 26.84 | 26.12 | 25.68 | Yes |
| 33. | 33 | 33 | M | POSITIVE | 27.47 | 26.84 | 26.22 | Yes |
| 34. | 34 | 38 | M | NEGATIVE | - | - |  | Yes |
| 35. | 35 | 44 | M | POSITIVE | 26.18 | 26.56 | 27.14 | **NO** |
| 36. | 36 | 45 | M | POSITIVE | 25.68 | 25.46 | 25.39 | **NO** |
| 37. | 37 | 39 | M | NEGATIVE | - | - |  | Yes |
| 38. | 38 | 26 | M | NEGATIVE | - | - |  | Yes |
| 39. | 39 | 30 | F | POSITIVE | 24.09 | 23.67 | 24.44 | Yes |
| 40. | 40 | 38 | M | NEGATIVE | - | - |  | Yes |
| 41. | 41 | 40 | F | NEGATIVE | - | - |  | Yes |
| 42. | 42 | 29 | M | NEGATIVE | - | - |  | Yes |
| 43. | 43 | 59 | F | NEGATIVE | - | - |  | Yes |
| 44. | 44 | 65 | F | POSITIVE | 24.66 | 24.46 | 24.02 | Yes |
| 45. | 45 | 73 | M | NEGATIVE | - | - |  | Yes |
| 46. | 46 | 54 | M | NEGATIVE | - | - |  | Yes |
| 47. | 47 | 42 | M | NEGATIVE | - | - |  | Yes |
| 48. | 48 | 39 | M | NEGATIVE | - | - |  | Yes |
| 49. | 49 | 28 | M | POSITIVE | 16.78 | 17.68 | 18.62 | Yes |
| 50. | 50 | 83 | F | POSITIVE | 28.07 | 27.84 | 26.83 | Yes |
| 51. | 51 | 58 | F | NEGATIVE |  |  |  | Yes |
| 52. | 52 | 61 | M | POSITIVE | 28.58 | 28.12 | 28.29 | Yes |
| 53. | 53 | 74 | F | NEGATIVE |  |  |  | Yes |
| 54. | 54 | 39 | M | POSITIVE | 26.78 | 26.14 | 26.98 | **NO** |
| 55. | 55 | 16 | F | POSITIVE | 26.19 | 27.04 | 26.87 | Yes |
| 56. | 56 | 19 | F | POSITIVE | 24.04 | 23.68 | 23.55 | Yes |
| 57. | 57 | 47 | M | NEGATIVE |  |  |  | Yes |
| 58. | 58 | 36 | F | POSITIVE | 17.88 | 18.12 | 18.36 | Yes |
| 59. | 59 | 25 | M | POSITIVE | 27.87 | 26.84 | 26.24 | Yes |
| 60. | 60 | 29 | F | NEGATIVE | - | - |  | Yes |
| 61. | 61 | 23 | M | POSITIVE | 23.94 | 23.66 | 24.15 | Yes |
| 62. | 62 | 48 | F | POSITIVE | 22 | 22.68 | 22.34 | Yes |
| 63. | 63 | 39 | M | NEGATIVE | - | - |  | Yes |
| 64. | 64 | 29 | M | POSITIVE | 25.46 | 24.48 | 22.48 | Yes |
| 65. | 65 | 51 | M | NEGATIVE | - | - |  | Yes |
| 66. | 66 | 52 | M | NEGATIVE | - | - |  | Yes |
| 67. | 67 | 48 | F | POSITIVE | 26.84 | 25.95 | 25.67 | Yes |
| 68. | 68 | 23 | F | NEGATIVE | - | - |  | Yes |
| 69. | 69 | 34 | F | NEGATIVE | - | - |  | Yes |
| 70. | 70 | 47 | M | POSITIVE | 28.16 | 27.68 | 28.49 | Yes |
| 71. | 71 | 46 | F | NEGATIVE | - | - |  | Yes |
| 72. | 72 | 48 | M | POSITIVE | 26.84 | 26.46 | 27.05 | Yes |
| 73. | 73 | 37 | F | NEGATIVE | - | - |  | Yes |
| 74. | 74 | 29 | M | NEGATIVE | - | - |  | **NO** |
| 75. | 75 | 58 | F | POSITIVE | 23.56 | 23.98 | 23 | Yes |
| 76. | 76 | 52 | M | POSITIVE | 30.27 | 29.61 | 29.98 | Yes |
| 77. | 77 | 65 | F | NEGATIVE |  |  |  | Yes |
| 78. | 78 | 61 | M | POSITIVE | 30.18 | 30.45 | 30.68 | Yes |
| 79. | 79 | 71 | M | NEGATIVE | - | - |  | Yes |
| 80. | 80 | 47 | M | NEGATIVE | - | - |  | **NO** |
| 81. | 81 | 27 | F | NEGATIVE | - | - |  | Yes |
| 82. | 82 | 38 | F | POSITIVE | 28.18 | 27.84 | 27.99 | Yes |
| 83. | 83 | 47 | M | NEGATIVE | - | - |  | Yes |
| 84. | 84 | 39 | F | NEGATIVE | - | - |  | Yes |
| 85. | 85 | 19 | F | NEGATIVE | - | - |  | Yes |
| 86. | 86 | 42 | F | NEGATIVE | - | - |  | Yes |
| 87. | 87 | 48 | M | POSITIVE | 28.64 | 28.87 | 28.15 | Yes |
| 88. | 88 | 88 | M | POSITIVE | 26.84 | 26.38 | 26.78 | **NO** |
| 89. | 89 | 23 | M | POSITIVE | 28.42 | 28.03 | 28.56 | Yes |
| 90. | 90 | 72 | M | NEGATIVE | - | - |  | Yes |
| 91. | 91 | 67 | M | POSITIVE | 29.87 | 28.14 | 29.16 | Yes |
| 92. | 92 | 58 | F | POSITIVE | 26.73 | 26.06 | 26.44 | Yes |
| 93. | 93 | 34 | F | NEGATIVE | - | - |  | Yes |
| 94. | 94 | 35 | F | NEGATIVE | - | - |  | Yes |
| 95. | 95 | 32 | M | NEGATIVE | - | - |  | Yes |
| 96. | 96 | 48 | M | POSITIVE | 27.64 | 27.16 | 27.88 | Yes |
| 97. | 97 | 56 | F | NEGATIVE | - | - |  | Yes |
| 98. | 98 | 45 | F | NEGATIVE | - | - |  | Yes |
| 99. | 99 | 56 | M | POSITIVE | 25.49 | 25.12 | 25.34 | Yes |
| 100. | 100 | 34 | M | POSITIVE | 26.78 | 25.88 | 26.45 | Yes |

